# Threonine synthase *CoTHR4* is involved in infection-related morphogenesis during the pre-penetration stage in *Colletotrichum orbiculare*

**DOI:** 10.1101/543827

**Authors:** Ken Harata, Tetsuro Okuno

## Abstract

Upon recognition of host plants, *Colletotrichum orbiculare*, an anthracnose disease fungus of cucurbitaceous plants, initiates morphological differentiation, including conidial germination and appressorium formation on the cuticle layer. The series of infection processes of *C. orbiculare* requires enormous nutrient and energy, but the surface of the cucurbitaceous hosts is hardly nutrient-rich. Hence, *C. orbiculare* must exert tight management of its intracellular nutrients in order to properly induce infection-related morphogenesis. Here, we carried out a large-scale insertional mutagenesis screen using *Agrobacterium tumefaciens*-mediated transformation to identify novel genes involved in the pathogenicity of *C. orbiculare* and found that *CoTHR4*-encoded threonine synthase, a homolog of *Saccharomyces cerevisiae THR4*, is required for pathogenicity and conidiation in *C. orbiculare*. Threonine supplementation allowed the *cothr4* mutant to produce conidia to a level equivalent to that of the wild-type. The conidia produced from the threonine-treated *cothr4* mutant failed to germinate in the absence of threonine, but retained the ability to germinate and to form appressoria in the presence of threonine. However, the conidia produced from the threonine-treated *cothr4* mutant remained attenuated in pathogenicity on cucumber cotyledons even in the presence of threonine. Cytorrhysis assays revealed that appressoria of the *cothr4* mutant induced by exogenous threonine treatment showed low turgor generation. Taken together, these results showed that threonine synthase CoThr4 plays a pivotal role in infection-related morphogenesis during the pre-penetration stage of *C. orbiculare*.

## Introduction

A prerequisite for the successful infection of host plants by phytopathogenic fungi is control of the elaborate infection-related morphogenesis. *Colletotrichum orbiculare*, which causes severe anthracnose disease symptoms on cucurbitaceous plants, undergoes dynamic morphogenesis for the establishment of infection. Upon recognition of physical signals on the plant surface, conidia of *C. orbiculare* germinate and develop a specialized infection cell, called an appressorium, which functions to rupture the plant cell walls by a mechanical driving force using intracellular turgor pressure [1, 2]. To date, functional analysis of *C. orbiculare* mutants defective in infection-related morphogenesis has uncovered two representative signal transduction pathways respectively mediated by mitogen-activated protein kinase (MAPK) and cAMP-dependent protein kinase (PKA) [3]. These pathways are widely conserved in eukaryotes and play a pleiotropic role in intracellular events. MAPK Cmk1, a homologue of MAPK Fus3/Kss1 in *Saccharomyces cerevisiae*, is necessary for conidial germination and appressorium formation in *C. orbiculare* [4]. The cAMP-PKA signaling pathway is essential for conidial germination in *C. orbiculare* [5]. The infection-related morphogenesis regulated by these pathways is common to *M. oryzae*, *Botrytis cinerea*, *Fusarium graminearum*, and *Ustilago maydis*, which cause devastating diseases of cultivated crops [6–10]. Moreover, positive and negative regulators of these signaling pathways have been characterized, and a detailed signaling network for infection-related morphogenesis has been identified in *C. orbiculare* and *M. oryzae* [11, 12]. However, although intracellular signal transduction in response to environmental signals has been well documented, knowledge of the intracellular nutrient metabolism underlying the cellular differentiation processes in *C. orbiculare* is less understood.

*C. orbiculare* and *M. oryzae* must adapt to nutrient starvation conditions on the plant surface during the pre-penetration stage, which includes conidial germination and appressorium formation. Under adverse conditions, these fungi adopt an infection strategy that utilizes the limited nutrient sources of conidia. In *C. orbiculare*, *ATG8*-mediated nonselective autophagy contributes to conidial germination and appressorium formation, and *ATG26*-mediated pexophagy may be involved in the cell wall integrity of the appressorium [13]. Nonselective autophagy-related genes are also associated with infection-related morphogenesis in *M. oryzae* [14]. Thus, valuable energy sources supplied by recycling machinery are exploited in the pre-penetration stage of phytopathogenic fungi. In addition to this mechanism, appropriate regulation of amino acid metabolism is reported to be associated with infection-related morphogenesis in *M. oryzae*. Glutamine synthase *GLN1* gene expression induced by loss of the GATA transcription factor *ASD4* increases intracellular glutamine levels, leading to activation of the TOR signaling pathway as a negative factor for appressorium formation [15]. Carbamoyl phosphate synthase *MoCPA2* affects appressorium formation and penetration into the host plant by regulating arginine biosynthesis [16]. Moreover, the biosynthesis of methionine, valine, leucine and isoleucine plays critical roles in conidiation and pathogenicity, suggesting that exquisite control of amino acid biosynthesis contributes to the appropriate infection cycle of *M. oryzae* [17–19].

Amino acid threonine biosynthesized from aspartate via five enzymatic reactions is required for auxotrophy and sporulation in *S. cerevisiae* [20, 21]. Threonine synthase, which plays a crucial role in the final step of the threonine biosynthesis pathway, is widely conserved in bacteria, fungi and plants, but not in mammals, and it catalyzes the production of threonine from O-phosphohomoserine [22–24]. Threonine synthase has been enthusiastically analyzed in the human-pathogenic fungi *Cryptococcus neoformans* and *Candida albicans*, and could be a target for a novel antifungal drug [25, 26]. However, the relation between threonine synthase and the infection process in phytopathogenic fungi, including *C. orbiculare*, has not been elucidated.

In this study, we set out to identify novel genes related to conidiation and pathogenicity of *C. orbiculare* through a large-scale forward genetic screening using *Agrobacterium tumefaciens*-mediated transformation. Consequently, we identified threonine synthase *CoTHR4*, a homolog of *THR4* in *S. cerevisiae*. Phenotypic and cytological analysis of the *cothr4* mutant revealed that CoThr4 functions to initiate the dynamic morphogenesis of *C. orbiculare* during the pre-penetration stage by modulating intracellular threonine.

## Materials and Methods

### Fungal and bacterial strains, culture conditions, and genomic DNA blot analysis

The *Colletotrichum orbiculare* (Berk. & Mont.) Arx [syn. *C. lagenarium* (Pass.); Ellis & Halst.] strain 104-T (MAFF240422) was used as the wild-type. All *C. orbiculare* strains were cultured on PDA (3.9% [w/v] PDA; Nissui Pharma, Tokyo) or PDA containing L-threonine at concentrations ranging from 0.1 to 5 mM at 24 °C. *Escherichia coli* DH5α -competent cells cultured in Luria-Bertani (LB) medium [27] at 37 °C were used as the host for gene manipulation. Kanamycin (50 μg/ml) was added to the medium as needed. *A. tumefaciens* C58C1 cultured in LB medium at 28 °C was used to transform *C. orbiculare* as previously described [12]. Total DNA extraction from the mycelia of *C. orbiculare* and DNA blot analysis were performed as previously described.

### Fungal transformation

*Agrobacterium tumefaciens*-mediated transformation (AtMT) in *C. orbiculare* was performed as previously described [28]. Hygromycin-resistant transformants were selected on PDA containing 50 μg/ml of hygromycin B, 75 μg/ml of cefotaxime and 75 μg/ml of spectinomycin (Wako Chemicals, Osaka, Japan). Sulfonylurea-resistant transformants were selected on SD medium (0.67 g yeast nitrogen base without amino acids, 2 g glucose, and 15 g agar in 1 L distilled water) containing 4 μg/ml of chlorimuron-ethyl (Maruwa Biochemical, Tokyo), 75 μg/ml of cefotaxime and 75 μg/ml of spectinomycin.

### Screening of conidiation- and pathogenicity-deficient mutants and identification of mutated genes

To obtain conidiation and pathogenicity-deficient mutants from 6996 T-DNA insertional lines generated by AtMT, the transformants were incubated on PDA. After incubation at 24 °C for 5 days, we selected transformants that showed reduced conidiation in comparison with that of the wild-type by visual observation. Next, to evaluate pathogenicity in the selected transformants, conidial suspensions were adjusted to 5.0×10^4^, 1×10^5^ or 5.0×10^5^ conidia/ml in accordance with the amount of conidia, respectively, and were placed on cucumber cotyledons. The inoculated leaves were incubated in a humid box at 24°C with a 16 h photoperiod for 6 days. Genomic DNA fragments flanking the inserted T-DNA in selected mutants were amplified by thermal asymmetrical interlaced PCR (Tail PCR) with specific primers and sequenced [28].

### Measurement of hyphal growth

For the observation of fungal growth, a Cork borer was used to punch mycelial disks (0.4 cm in diameter) of each strain from the marginal region of the colony after 7 days of incubation, and the disks were placed onto SD medium with or without 1 mM threonine. After 7 days of incubation, the colony diameters were measured.

### Plasmid construction

All cloning for plasmid construction was carried out using an In-Fusion HD Cloning Kit (Clontech, Palo Alto, CA). All primers used in this study are listed in Table S1.

For the construction of the *CoTHR4* gene complementation plasmid (PBIG4MRSCoTHR4), an approximately 5.0 kb *CoTHR4* fragment, including the 1.5 kb region upstream from the *CoTHR4* start codon and 1.7 kb *CoTHR4* region downstream from the *CoTHR4* stop codon, and the pBIG4MRSrev vector fragment, including the sulfonylurea-resistance gene, were amplified by PCR with the primer pairs CoTHR4F1B/CoTHR4R1A and pBIG4MRBSF1A/pBIG4MRBSF1AR1A, respectively. PBIG4MRSCoTHR4 was generated by inserting amplified *CoTHR4* fragments into the linearized pBIG4MRSrev vector.

For the construction of the *cothr4* gene-replacement plasmid (pBIG4MRScothr4), an approximately 1.4 kb fragment of the hygromycin-resistance gene (HPH) and a pBIG4MRSCoTHR4 vector fragment not including the *CoTHR4* ORF region were amplified by PCR with the primer pairs HPHF1B/HPHR1A and pBIcothr4F1A/pBIcothr4R1B, respectively. pBIG4MRScothr4 was generated by inserting HPH into the linearized pBIG4MRSCoTHR4 vector.

For the construction of the *CoTHR4*-*mCherry* fusion gene plasmid (pBIG4MRSCoTHR4mC), the mCherry fluorescent gene fragment (mCherry) and the pBIG4MRSCoTHR4 vector fragment were amplified by PCR using the primer pairs mCherryF1B/mCherryR1A and pBICoTHR4mCF1A/pBICoTHR4mCR1B, respectively. pBIG4MRSCoTHR4mC was generated by inserting the mCherry gene into the linearized pBIG4MRSCoTHR4 vector.

### Pathogenicity assay

Pathogenicity assays were performed using conidia suspended in distilled water containing threonine at different concentrations. Ten μl conidial suspensions (5.0×10^4^ conidia/ml) were placed on 6 spots per detached cucumber cotyledon, and inoculated leaves were incubated in a humid box at 24°C with a 16 h photoperiod for 6 days.

### Cytorrhysis assay

For evaluation of turgor pressure within the appressorium, conidia of each strain were suspended in 1 mM threonine solution, and 20 μl conidial suspensions (5.0×10^4^ conidia/ml) were placed on three spots per coverslip. After 24 h of incubation, each sample was exposed to glycerol or PEG 6000 solution for 15 min.

### Microscopy observation

For observation of infection-related morphogenesis, conidia of each strain were suspended in distilled water, 0.1 mM or 1 mM threonine solution, and 20 μl conidial suspensions (5.0× 10^4^ conidia/ml) were placed on eight spots per coverslip. After 24 h of incubation, the conidial germination and the appressoria formation of each strain were observed using For observation of CoThr4:mCherry localization during appressorium development and LifeAct-RFP localization in the matured appressoria, 20 μl conidial suspensions (1.0×10^5^ conidia/ml) were placed and incubated on the coverslip and the abaxial surface of cucumber cotyledons. mCherry and RFP fluorescence signals were observed using a Leica SP8 confocal laser scanning microscope equipped with a diode-pumped solid state 552 nm laser; mCherry and RFP signals were detected from 600–630 nm and 590–630 nm by using an SP8 hybrid detector, respectively. All images were taken using a 63×oil immersion lens.

## Results

### Screening for mutants deficient in conidiation and pathogenicity

We generated 6996 transformants through large-scale insertional mutagenesis using *Agrobacterium tumefaciens*-mediated transformation (AtMT). As a primary screening, we selected the mutants whose colonies appeared to have a reduced amount of conidia by visual observation, and obtained 147 independent mutants. We then carried out pathogenicity assays on cucumber cotyledons using 136 of these mutants, because the amount of conidia was insufficient to perform the pathogenicity test in 11 mutants. Finally we obtained 40 independent mutants that showed an attenuated pathogenesis relative to the wild-type after two rounds of pathogenicity assays on cucumber cotyledons. Genomic DNA segments adjacent to T-DNA insertion sites in the selected mutants were isolated by thermal asymmetrical interlaced-polymerase chain reaction and the amplified products were sequenced. Eventually, we determined T-DNA insertion sites in 17 mutant lines and identified 18 genes as candidates for involvement in conidiation and pathogenicity (Table 1). In this paper, we further investigated one of the 17 mutant lines-namely, CPD1 (<u>c</u>onidiation and <u>p</u>athogenesis <u>d</u>eficient mutant).

**Table 1.** T-DNA insertion site of conidiation and pathogenicity defective mutant in *C. orbiculare*.

| T-DNA insertion line | Putative T-DNA insetion site | Blast hit description |
| --- | --- | --- |
| CPD1 | Cob_04177:ORF | threonine synthase |
| CPD2 | Cob_04238:ORF | Hym1 protein |
| CPD3 | Cob_11154:Upstream 194bp | transformer-sr ribonucleoprotein |
| CPD4 | Cob_13036:ORF | nascent polypeptide-associated complex subunit alpha |
| CPD5 | Cob_04426:ORF | kinase activator |
| CPD6 | Cob_07204:Upstream432bp | arsenate reductase |
|  | Cob_07205:upstream367bp | hypothetical protein |
| CPD7 | Cob_04884:ORF | ell-associated factor |
| CPD8 | Cob_09403:Upstream 142bp | c-x8-c-x5-c-x3-h type zinc finger protein |
| CPD9 | Cob_10557:ORF | peroxisome biosynthesis protein (pas10 peroxin-12) |
| CPD10 | Cob_12838:Downstream 841bp | phd and ring finger domain-containing protein c |
|  | Cob_12839:Upstream 219bp | 50s ribosomal subunit l30 |
| CPD11 | Cob_07968:ORF | cral trio domain protein |
| CPD12 | Cob_00044:ORF | nadh-ubiquinone oxidoreductase subunit grim-19 |
| CPD13 | Cob_04238:ORF | Hym1 protein |
| CPD14 | Cob_09202:ORF | serine threonine protein kinase |
| CPD15 | Cob_05343:Upstream 2233bp | sh3 domain containing protein |
| CPD16 | Cob_09249:ORF | hytotehtical protein |
| CPD17 | Cob_05102:Upstream 1018bp | pq loop repeat protein |

### CPD1 has a mutation in the threonine synthase gene

CPD1 showed a decrease in conidiation, and the conidia caused much smaller lesions on the cucumber cotyledons compared with the large severe lesions induced by the wild-type (S1A-C Fig). The T-DNA was inserted into the 204-bp region downstream from the start codon of Cob_04177 in CPD1 (S1D Fig). A BLASTp search indicated that the amino acid sequences deduced from this gene shared 54% identify with that from threonine synthase *THR4* of *S. cerevisiae* (S2 Fig). Therefore, we name the gene Cob_04177 as *CoTHR4*, and we subjected this gene to functional analysis.

### Threonine synthase *CoTHR4* is required for conidiation and pathogenicity

To perform functional analysis of the *CoTHR4* gene, we generated a *cothr4* disruption mutant by double crossover homologous recombination using AtMT. Disruption of the targeted gene was confirmed by Southern blotting (S3 Fig). The colony of the *cothr4* mutant was smaller than that of the wild-type on potato dextrose agar (PDA) medium (Fig 1A). Conidia production was dramatically reduced in the *cothr4* mutant and the *CoTHR4*-complemented transformant produced an equivalent number of conidia as the wild-type, indicating that *CoTHR4* plays a critical role in conidiation (Fig 1B). Next, to examine the requirement of *CoTHR4* for fungal pathogenicity, we inoculated cucumber cotyledons with conidia of the *cothr4* mutant. At 6 days postinoculation (dpi), the wild-type and the *CoTHR4*-complemented transformant induced severe disease symptoms on inoculation sites, whereas the *cothr4* mutant was unable to cause lesions, suggesting that *CoTHR4* is essential for the pathogenicity (Fig 1C).

**Fig 1.**
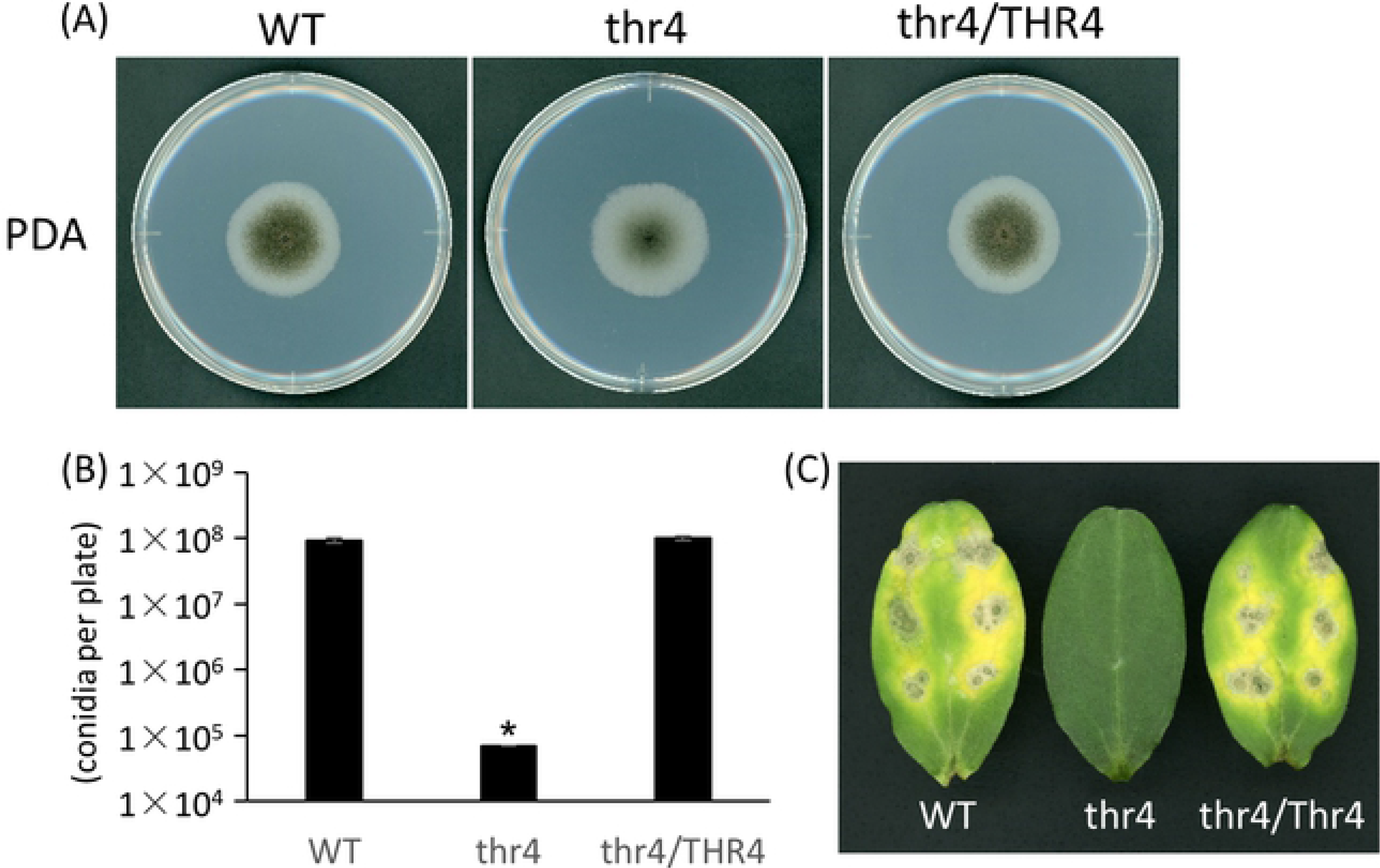
*CoTHR4* is required for conidiation and pathogenesis in *C. orbiculare*. **(A)** Colony phenotype and hyphal growth of the *cothr4* mutant on PDA medium. A mycelial block of the wild-type (WT), the *cothr4* mutant (thr4), and *CoTHR4*-complemented transformant (thr4/THR4) was placed on PDA medium and incubated at 24 °C for 5 days. **(B)** The average number of conidia in a colony of WT, thr4, and thr4/THR4 on the PDA medium after 5 days incubation at 24 °C. Error bars represent the means of standard errors (n = 5). Asterisk represents significant difference between the wild-type and the *cothr4* mutant (Student’s *t* test: *P < 0.01). **(C)** Pathogenicity assay of the *cothr4* mutant on the detached cucumber cotyledons. Cucumber cotyledons were drop-inoculated with conidial suspension of WT, thr4, and thr4/THR4, and incubated at 24 °C for 6 days.

### Defects in hyphal growth and conidiation of the *cothr4* mutant were fully restored by exogenously supplied threonine

Amino acids, such as arginine, methionine, leucine, isoleucine and valine, are essential for hyphal growth in phytopathogenic fungi [17–19, 29]. To examine whether threonine is necessary for fungal viability, we observed hyphal growth of the *cothr4* mutant on the synthetic defined (SD) media with or without threonine. Hyphal growth of the *cothr4* mutant was completely abolished in the absence of threonine, whereas colony growth of the *cothr4* mutant was similar to that of the wild-type and *CoTHR4*-complemented transformant in SD media supplemented with 1 mM threonine (Fig 2A and 2B). These results indicated that threonine is essential for hyphal growth of the *cothr4* mutant.

**Fig 2.**
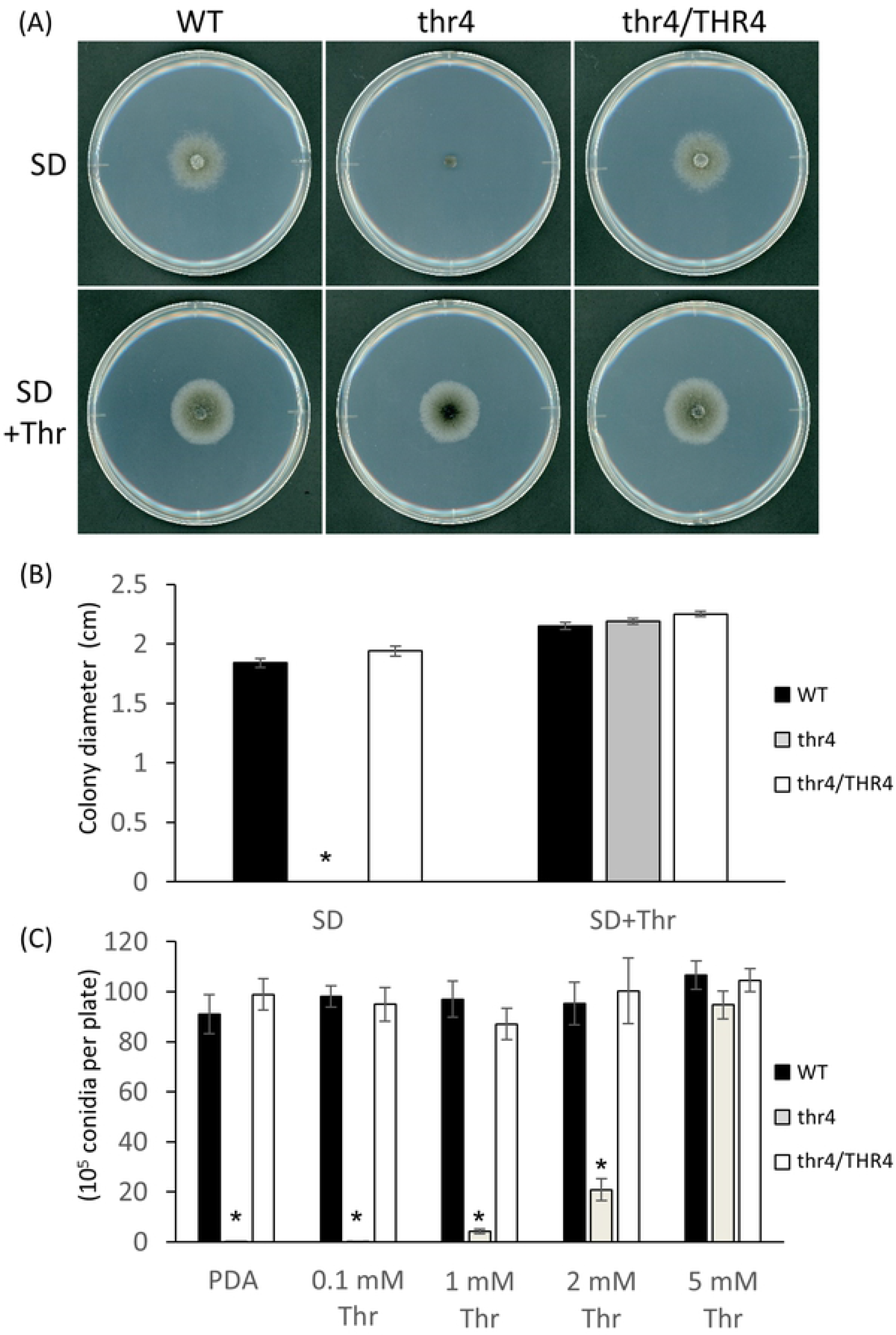
Hyphal growth and conidiation of the *cothr4* mutant require threonine. **(A)** Hyphal growth of the *cothr4* mutant on SD medium supplemented with threonine. A mycelial block of each strain was placed on SD medium with or without 1 mM threonine and incubated at 24 °C for 7 days. SD; SD medium, SD+Thr; SD medium with 1 mM threonine. **(B)** The average size of a colony diameter in the *cothr4* mutant on the SD medium supplemented with threonine. Error bars represent the means of standard errors (n = 5). Asterisk represents significant difference between the wild-type and the *cothr4* mutant (Student’s *t* test: *P < 0.01). **(C)** The average number of conidia in a colony of the *cothr4* mutant on the PDA medium supplemented with 0.1, 1, 2, or 5 mM threonine after 5 days incubation at 24 °C. Error bars represent the means of standard errors (n = 5). Asterisks represent significant difference between the wild-type and the *cothr4* mutant (Student’s *t* test: *P < 0.01).

Because the *cothr4* mutant produced only a few conidia on the PDA medium (Fig 1B), we next investigated whether the production of conidia in the *cothr4* mutant would be stimulated by exogenously supplied threonine. We measured the amount of conidia produced from the *cothr4* mutant grown on the PDA medium supplemented with threonine at different concentrations ranging from 0.1 mM to 5 mM. The amount of conidia produced from the *cothr4* mutant increased as the concentration of supplemented threonine increased, and reached the wild-type levels in 5 mM threonine-supplemented PDA media (Fig 2C). The wild-type and *CoTHR4*-complemented transformant produced abundant conidia (>8×10^5^ conidia per colony) in all threonine-supplemented PDA media (Fig 2C). This result also showed that threonine in concentrations up to at least 5 mM has no toxic effect on conidiation in *C. orbiculare*. Taken together, these results suggested that threonine is important for conidiation in *C. orbiculare*.

### Exogenously supplied threonine stimulated conidial germination and appressorium formation in the *cothr4* mutant

*CoTHR4* could be involved in any steps of infection-related morphogenesis, because the conidia obtained from the *cothr4* mutant failed to cause visible lesions on cucumber leaves (Fig 1C). First, we tested the ability of the mutant to germinate and form appressorium on coverslips in the absence of threonine using the conidia of the *cothr4* mutant produced on PDA media supplemented with different concentrations of threonine (see Fig 2C). More than 90% of conidia of the wild-type and *CoTHR4*-complemented transformant produced on all threonine-supplemented PDA media germinated and formed appressoria, whereas more than 90% of conidia of the *cothr4* mutant failed to germinate, except for those obtained from 1 mM threonine-supplemented PDA media, whose germination rate was approximately 25% (Fig 3). These results suggested that threonine biosynthesis is required for efficient germination in conidia. To observe the expression and localization of CoThr4 in conidia before germination, we generated the *CoTHR4*:*mCherry* construct, which expresses the C-terminally *mCherry* gene-fused *CoTHR4* under its native promoter, and used it to transform the *cothr4* mutant. The *CoTHR4*:*mCherry*-introduced transformant (CoTHR4:mCherryCom) restored full pathogenesis on cucumber cotyledons, indicating that *CoTHR4*:*mCherry* is functional (S4 Fig). CoThr4:mCherry fluorescence was observed as small localized punctuates in ungerminated conidia of CoTHR4:mCherryCom on a coverslips at 15 min and at 2 h after incubation, while its signals were much weaker or not detectable in the germinated conidia and in the appressorium developed from the germ tube tip of conidia (S5 Fig). These results suggested that threonine was synthesized and accumulated in ungerminated conidia.

**Fig 3.**
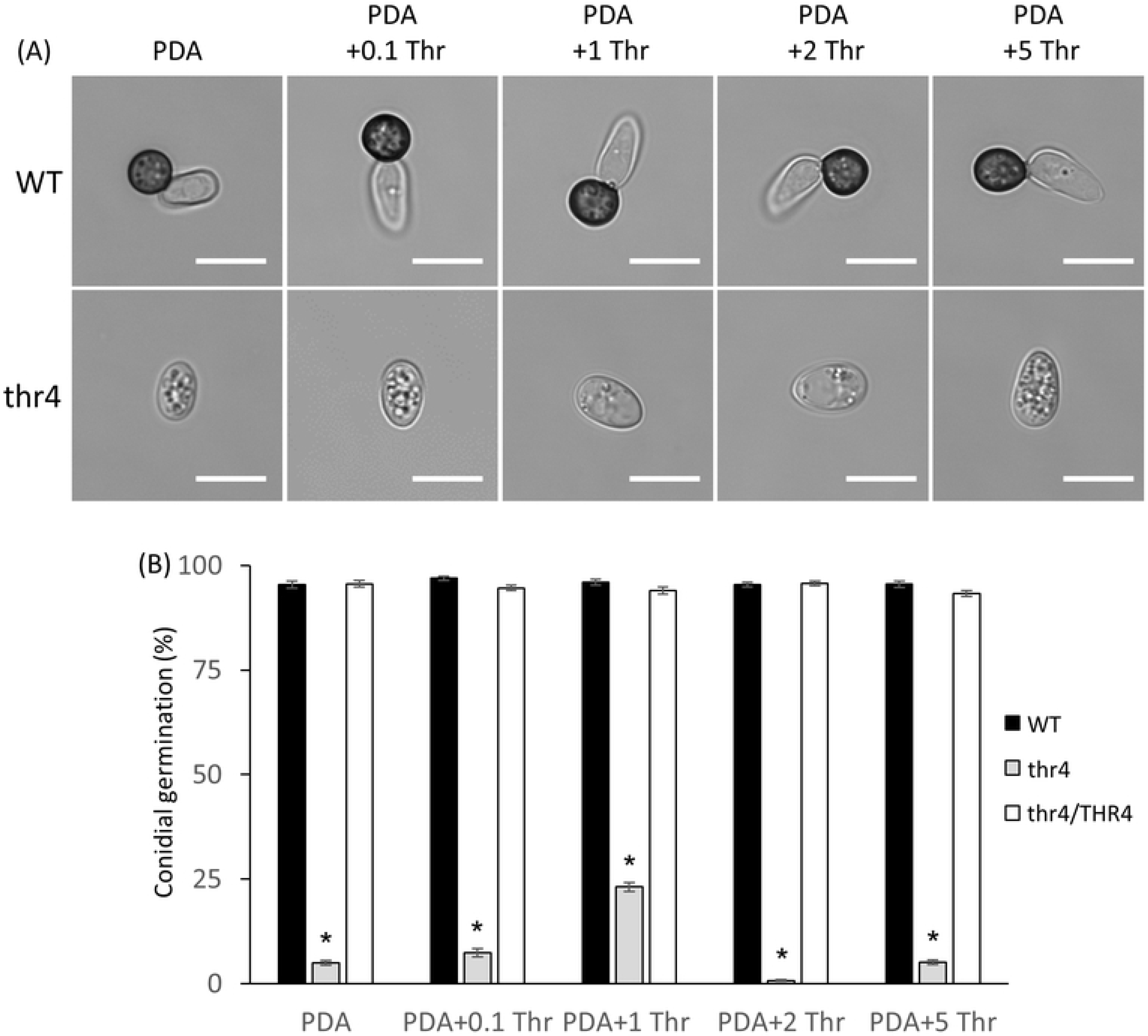
Conidia of the threonine-treated *cothr4* mutant were unable to germinate in the absence of threonine. **(A)** Appressorium differentiation of conidia of the wild-type and the *cothr4* mutant produced on PDA medium supplemented with threonine at different concentrations ranging from 0.1 mM to 5 mM on coverslip at 24 h after incubation. PDA; PDA medium, PDA+0.1 Thr; PDA medium supplemented with 0.1 mM threonine, PDA+1 Thr; PDA medium supplemented with 1 mM threonine, PDA+2 Thr; PDA medium supplemented with 2 mM threonine, PDA+5 Thr; PDA medium supplemented with 5 mM threonine. Scale bar, 10 μm. **(B)** The frequency of germination in conidia of the *cothr4* mutant. Approximately 100 conidia of each strain were observed per spot, and eight spots were examined. Three independent experiments were conducted, and standard errors were indicated. Asterisks represent significant difference between the wild-type and the *cothr4* mutant (Student’s *t* test: *P < 0.01).

Next, we tested whether exogenously supplied threonine could stimulate infection-related morphogenesis in the conidia of the *cothr4* mutant produced on 5 mM threonine-supplemented PDA. More than 85% of conidia of the *cothr4* mutant germinated and formed appressoria in the presence of 0.1 mM or 1 mM threonine at frequencies similar to those of the wild-type and *CoTHR4*-complemented transformant. Taken together, these results suggested that threonine plays an important role in triggering initiation of conidial germination followed by appressorium formation (Fig 4).

**Fig 4.**
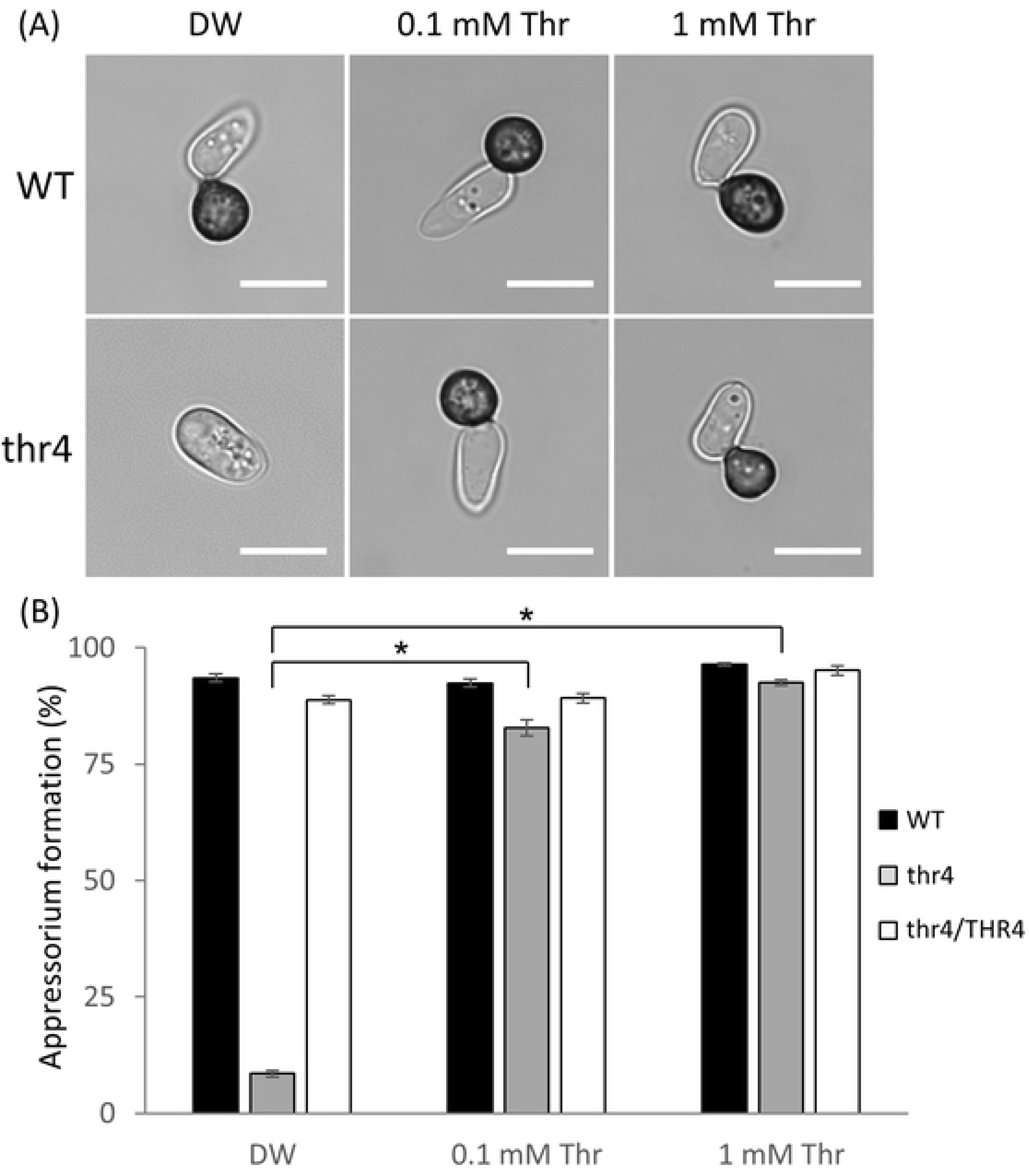
Conidia produced from the threonine-treated *cothr4* mutant induced appressorium formation in the presence of threonine. Conidia of each strain produced on the 5 mM threonine-supplemented PDA medium were suspended by distilled water, 0.1, or 1 mM threonine solution and incubated on coverslips at 24 °C for 24 h. DW; distilled water, 0.1 mM Thr; 0.1 mM threonine solution, 1 mM Thr; 1 mM threonine solution. **(A)** Appressorium formation of the *cothr4* mutant. Scale bar, 10 μm. **(B)** The frequency of appressorium formation of the *cothr4* mutant. Approximately 100 conidia of each strain were observed per spot, and eight spots were examined. Three independent experiments were conducted, and standard errors were indicated. Asterisks represent significant difference between threonine-untreated each strain and threonine-treated each strain (Student’s *t* test: *P < 0.01).

### Conidia produced from the threonine-treated *cothr4* mutant were impaired in appressorium-mediated infection on cucumber cotyledons even in the presence of threonine

To examine whether the conidia produced from the threonine-treated *cothr4* mutant have the ability to cause disease on the host plants, the conidia suspended in distilled water containing 0.1 mM or 1 mM threonine were inoculated onto cucumber cotyledons and the leaves were incubated for 6 days. The conidia from the threonine-treated *cothr4* mutant had no or weak ability to cause disease symptoms even in the presence of threonine (Fig 5A). The results suggested that the *cothr4* mutant might be impaired in appressorium-mediated penetration or invasive growth in the epidermal cells, because the conidia produced from the threonine-treated *cothr4* mutant were able to develop appressoria in the presence of threonine. The observation of infection hyphae at 3 days after inoculation of cucumber leaves showed that only a few appressoria developed infection hyphae in the *cothr4* mutant, unlike in the wild-type and *CoTHR4*-complemented transformant (Fig 5B and 5C). Interestingly, conidia from the threonine-treated *cothr4* mutant formed appressorium-mediated penetration hyphae on cellulose membranes in the presence of threonine (S6 Fig). We considered that the pathogenicity defect of the *cothr4* mutant is due to impairment of the appressorium-mediated infection machinery on cucumber leaves.

**Fig 5.**
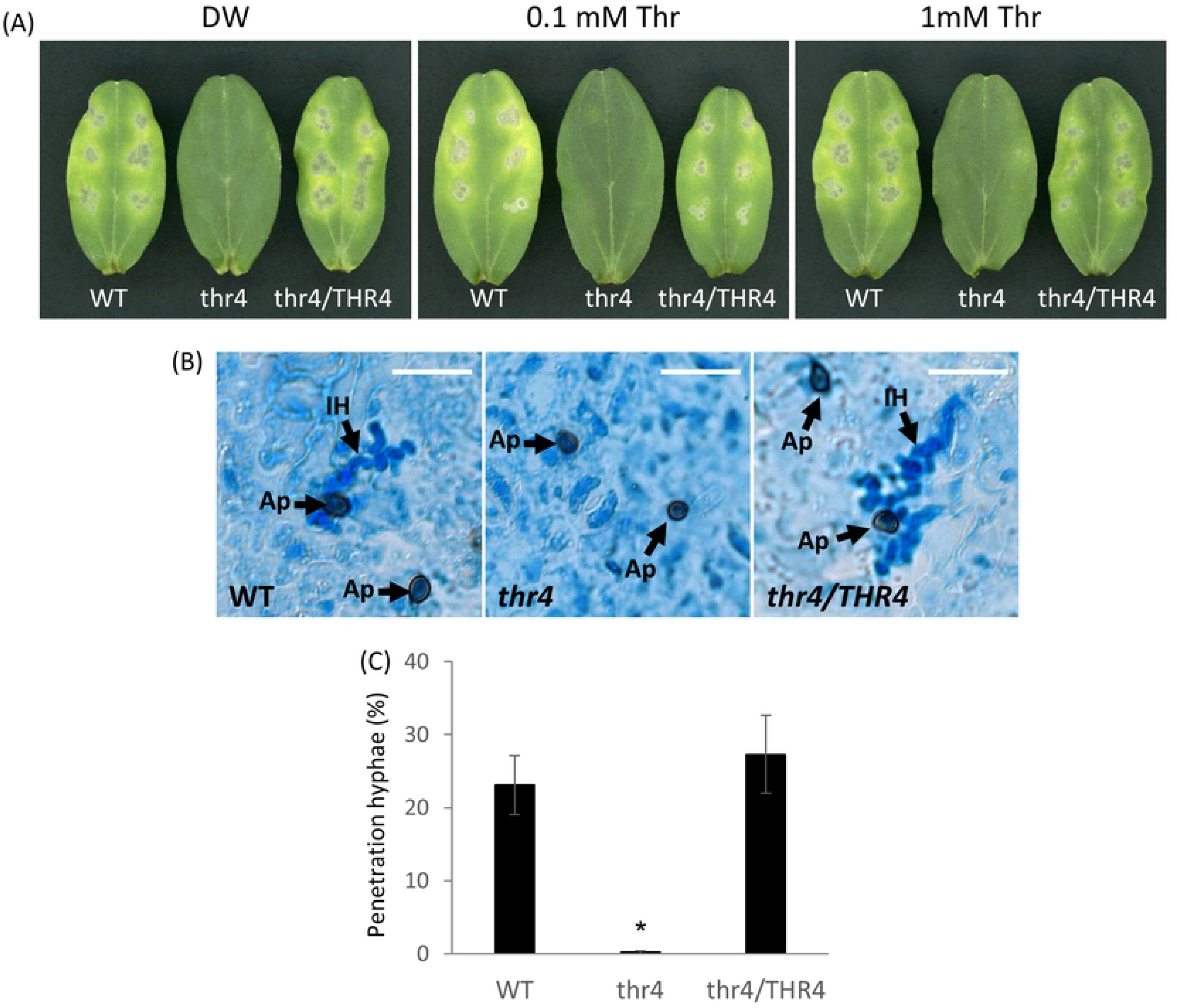
Conidia produced from the threonine-treated *cothr4* mutant were impaired in appressorium-mediated penetration on the cucumber leaves in the presence of threonine. Conidia of each strain produced on 5 mM threonine-supplemented PDA medium were suspended by distilled water, 0.1 mM or 1mM threonine solution. Conidial suspensions of each strain were placed on adaxial surface (A) or abaxial surface (B, C) of cucumber cotyledons. **(A)** Pathogenicity assays of the *cothr4* mutant on the cucumber cotyledons. **(B)** Infection hyphae development of the *cothr4* mutant. Conidia of each strain were suspended by 1 mM threonine and the inoculated leaves were incubated at 24 °C for 3 days. Ap; Appressorium, IH; Invasive Hyphae. Scale bar; 20 μm. **(C)** The frequency of the penetration hyphae of the *cothr4* mutant on the abaxial surface of cucumber cotyledons. Approximately 100 appressoria were observed per inoculation site, and three replications were examined. Three independent experiments were conducted, and standard errors were indicated. Asterisks represent significant difference between the wild-type and the *cothr4* mutant (Student’s *t* test: *P < 0.01).

The appressorium-mediated infection requires high turgor generation and cytoskeletal remodeling within the appressorium. Turgor pressure provides the mechanical force for penetration into the host cuticle and cell wall. We evaluated turgor pressure in the appressoria of the *cothr4* mutant using a cytorrhysis assay with glycerol. Interestingly, the frequency of appressorium collapse was significantly lower in the *cothr4* mutant than in the wild-type over any range of glycerol concentrations (Fig 6A). In *M. oryzae,* an *alb1* mutant that is defective in melanin biosynthesis, which is necessary for turgor generation, develops inappropriate porosity in the appressorial cell wall and thereby allows the inflow of glycerol, resulting in quick reinflation of the collapsed cell [30]. Consistent with the report of an albino mutant in *M. oryzae* [30], the appressoria of the melanin-deficient mutant (*pks1*) in *C. orbiculare* remained intact in the cytorrhysis assay using glycerol (Fig 6A). Therefore, we conducted a cytorrhysis assay using different concentrations of polyethylene glycol 6000 (PEG 6000), which is larger than glycerol in molecular size, and monitored the collapsed appressoria. The frequency of appressorium collapse was higher in the *cothr4* mutant than in the wild-type following 30% and 35% PEG 6000 treatment (Fig 6B). In the 40% PEG treatment, there was no difference in the frequency of appressorium collapse between the wild-type and the *cothr4* mutant (Fig 6B). These results suggested that the appressorial turgor generation of the *cothr4* mutant induced by exogenous threonine treatment did not reach a level comparable to that in the wild-type.

**Fig 6.**
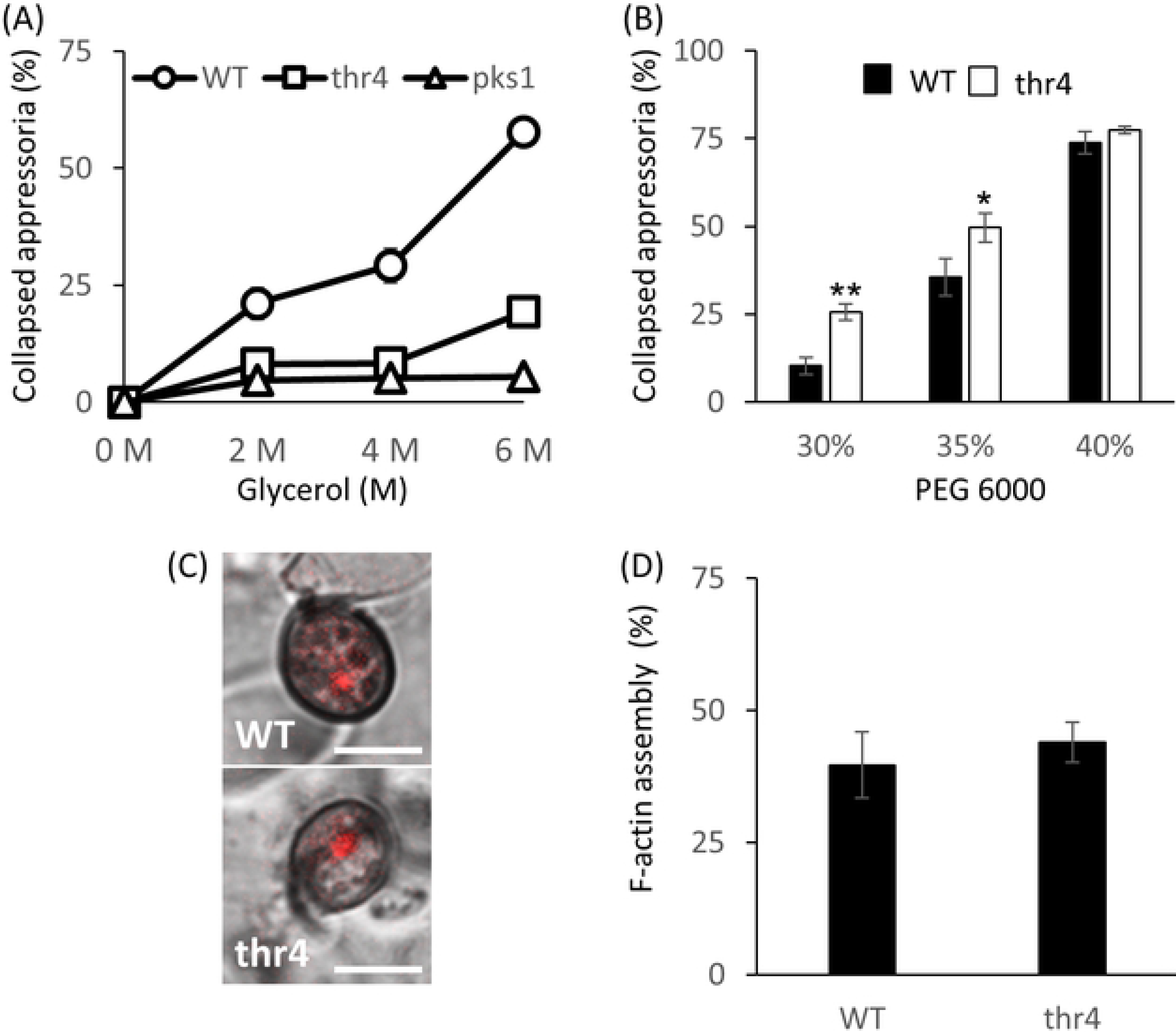
Conidia produced from the threonine-treated *cothr4* mutant showed a defect of appressorial turgor generation in the presence of threonine. **(A)** The frequency of appressorium collapse in the *cothr4* mutant exposed to glycerol at different concentrations ranging from 0 M to 6M. Conidia of each strain were suspended by 1 mM threonine and incubated on coverslips at 24 °C for 24 h. Approximately 100 appressoria were observed per spot, and three spots were examined. Three independent experiments were conducted, and standard errors were indicated. **(B)** The frequency of appressorium collapse in the *cothr4* mutant exposed to PEG 6000 solutions ranging from 30% to 40%. Asterisks represent significant difference between the wild-type and the *cothr4* mutant (Student’s *t* test: *P < 0.05, **P < 0.01). **(C)** LifeAct-RFP fluorescence in appressoria of the *cothr4* mutant on the abaxial surface of cucumber cotyledons. Conidia of each strain were suspended by 1 mM threonine and the inoculated leaves were incubated at 24 °C for 3 days. Scale bar; 5 μm. **(D)** The frequency of appressoria with a fluorescent signal of punctate structures in the *cothr4* mutant expressing LifeAct-RFP at 48 h postinoculation. Accumulation of LifeAct-RFP fluorescence in appressorium was examined and approximately 100 appressoria were observed. Three independent experiments were conducted, and standard errors were indicated.

The cytoskeletal remodeling event that assembles F-actin rings at the base of the appressorium is essential for organization of the appressorium pore where the penetration peg emerges [31–33]. To monitor F-actin organization in the appressorium of the *cothr4* mutant, we expressed the actin filament marker Lifeact-RFP in the *cothr4* mutant and the wild-type, and observed RFP signals within the appressorium before emergence of the penetration peg. In the wild-type, RFP fluorescence was localized to the punctate structures, which are envisaged as an F-actin assembly at the base of the appressorium (Fig 6C). Similar localization patterns were also detected in the *cothr4* mutant, and the frequency of appressoria accompanied with F-actin assembly signals was equivalent to that of wild-type (Fig 6C and 6D). These results suggested that *CoTHR4* has no effect on the cytoskeletal remodeling of appressorium pores for penetration peg formation.

## Discussion

Forward genetic screening using transformants generated by AtMT is a powerful approach for identifying pathogenicity genes in an unbiased manner, and various genes related to pathogenicity have been characterized in *Colletotrichum* species and *M. oryzae* [28, 34, 35]. In this study, we identified 18 candidates as pathogenicity and conidiation-related genes and dissected the role of threonine synthase *CoTHR4* in the life cycle of *C. orbiculare*, ranging from conidial development to appressorium development followed by infection peg formation and invasive growth. The characterization of auxotrophy mutants lacking amino acid biosynthesis in phytopathogenic fungi has revealed the diverse effects of amino acids on the infection processes. Here, we presented the first evidence that threonine synthesized by threonine synthase and threonine itself are important for the efficient conidiation and induction of infection-related morphogenesis in *C. orbiculare* and for its pathogenicity.

It is known that multiple amino acids, such as arginine, isoleucine, leucine and methionine, are required for conidial development in *M. oryzae* and *Fusarium graminearum* [17–19, 29]. Genome-wide gene expression profiling during conidiation in *M. oryzae* revealed the induction of many genes involved in amino acid transport [36]. Thus, several amino acids are involved in the production of conidia from mycelia in phytopathogenic fungi. Although arginine, isoleucine, leucine and methionine biosynthesis-related genes are indispensable for conidial development in *M. oryzae*, these genes are dispensable for conidial germination and appressorium formation in the fungus [17, 19, 37]. Interestingly, in contrast to the context of amino acid biosynthesis genes in *M. oryzae*, the majority of conidia of the *cothr4* mutant produced on the threonine-supplemented PDA media were unable to germinate and the conidia required exogenously supplied threonine for efficient germination. Regulation of conidial germination in *C. orbiculare* is known to involve the Ras/MAPK and cAMP pathways, which serve in the transmission of environmental signals to intracellular systems for initiation of infection-related morphogenesis, but how cellular nutrients facilitate the initial infection processes remains elusive [12]. Our results suggested that threonine production regulated by threonine synthase in conidia could potentially trigger conidial germination by functioning as a cellular nutrient signal. In *M. oryzae*, elevated levels of intracellular glucose and glutamine disturb appressorium development by activating the TOR pathway, which plays a pivotal role as an intracellular nutrient sensor [15, 38]. However, conidial germination in *M. oryzae* and *C. orbiculare* is unaffected in the presence of the TOR inhibitor rapamycin, suggesting that TOR activation is dispensable for conidial germination [15, 39]. Conidial germination might require a threonine-responsive system rather than the TOR pathway in *C. orbiculare*.

Hydrophobic surfaces function as a physical signal to effectively induce conidial germination in *orbiculare* [40]. This germination mechanism driven by environmental cues is conserved among filamentous fungi [41–43]. We found that conidia of the threonine-treated *cothr4* mutant were unable to germinate on a hydrophobic surface. According to proteomic analysis in *M. oryzae,* MoRgs proteins, which act as a negative regulator of G-protein during the infection-related morphogenesis in response to hydrophobic surfaces, have the potential to promote amino acid metabolism [44, 45]. Although there was no apparent interaction between threonine and intracellular signal transduction factors in response to external signals, there is a possibility that cellular threonine status might be controlled by a signal transduction factor that is instigated by the recognition of environmental signals, thereby leading to induction of conidial germination.

The conidia produced by the threonine-treated *cothr4* mutant germinated and formed appressoria in the presence of threonine, but failed to form infection hyphae (Fig 5B and 5C). The generation of turgor pressure in the appressorium is a prerequisite to proceed into an entry phase, and the tuning of this cellular event allows breakdown of the plant cell barrier [46, 47]. Cytological observations showed that appressoria of the *cothr4* mutant induced by exogenously supplied threonine exhibited a reduction in turgor generation. In *C. orbiculare* and *M. oryzae*, the melanin layer in the appressorial cell wall is a semipermeable membrane that blocks the efflux of glycerol from the appressorium, leading to high turgor pressure [47, 48]. We noticed that melanization in appressoria of the *cothr4* mutant induced by exogenously supplied threonine was slightly weaker than that in the wild-type (Fig 4). We thus considered that the decrease of turgor generation in appressoria of the *cothr4* mutant could be due to inadequate melanization in the appressorium, although ultrastructural analysis of the appressorial cell wall will be required to show an association between the two effects.

As described herein, the *cothr4* mutant requires an exogenous supply of threonine for efficient conidiation and appressorium formation. Interestingly, the appressoria of threonine-treated *cothr4* mutant conidia were non-functional in the presence of threonine. It should be noted that wild-type conidia developed functional appressoria and caused lesions on cucumber cotyledons in the presence of threonine at the concentrations used in the assays for the *cothr4* mutant conidia (Fig 5). These results raised the question of how CoThr4 participates in the development of functional appressoria. In *S. cerevisiae* and *C. albicans,* appropriate threonine biosynthesis is controlled by feedback regulation of aspartate kinase Hom3 and homoserine kinase Thr1 in the threonine biosynthetic pathway in response to elevated levels of threonine [26]. This mechanism is also conserved in *Arabidopsis thaliana*, and thus a threonine-sensing mechanism is thought to maintain cellular homeostasis [23]. Therefore, we assumed that, upon recognizing an abundance of threonine, CoThr4 might serve as a negative regulator of threonine biosynthesis or a coordinator of cellular homeostasis required for the appressorium-mediated infection.

## Acknowledgements

We are grateful to Dr. Yasuyuki Kubo for providing *Agrobacterium tumefaciens* strain C58C1 and pBIG4MRSrev plasmid, and Dr. Yoshitaka Takano for providing *pks1* mutant.

## Supporting information

**S1 Fig. Phenotype of CPD1 mutant.**

**(A)** Hyphal growth of CPD1 on PDA medium. A mycelial block of each strain was placed on PDA medium and incubated at 24 °C for 7 days.

**(B)** Pathogenicity assay of CPD1 on cucumber cotyledons. Conidial suspensions of each strain were placed on detached cotyledons of cucumber and incubated at 24 °C for 6 days.

**(C)** The average number of conidia in a colony of WT and CPD1 on the PDA medium at 24 °C after 5 days incubation. Error bars represent the means of standard errors (n = 5). Asterisk represents significant difference between the wild-type and CPD1 (Student’s *t* test: *P < 0.01).

**(D)** Schematic diagram of T-DNA insertion site in CPD1.

**S2 Fig. Amino acid sequence alignment of *C. orbiculare* CoThr4 with *S. cerevisiae* Thr4.**

Amino acids of *C. orbiculare* CoThr4 were aligned with those of *S. cerevisiae* Thr4 using the Clustal W program and shaded using Gene Doc. Numbers on the right indicate amino acid residue positions. Identical amino acids are indicated by a black background. Gaps introduced for alignments are indicated by a hyphen.

**S3 Fig. Gene disruption of *CoTHR4* in *C. orbiculare.***

**(A)** Schematic diagram of *CoTHR4* gene disruption construct in *C. orbiculare* by *Agrobacterium tumefaciens-*mediated transformation with the *cothr4* disruption vector to replace hygromycin phosphotransferase gene (hph) fragment with the *CoTHR4* gene. Bars represent probes for DNA gel blot. Following double crossover homologous recombination, a *Eco*RV fragment of approximately 1.8 kb containing *CoTHR4* in the wild type is predicted to be replaced by a fragment of approximately 5.9 kb containing the hph fragment.

**(B)** *CoTHR4* gene disruption was confirmed by Southern blot analysis. Genomic DNAs from the wild-type 104-T and transformants were digested with *Eco*RV and probe with an upstream 1.0 kb fragment of the *CoTHR4* gene. WT, wild-type 104-T; dis1-2, *cothr4* mutant.

**S4 Fig. Pathogenicity assay of *CoTHR4*:*mCherry*-introduced transformant on cucumber cotyledons.**

The inoculated cucumber cotyledons were incubated at 24 °C for 6 days. WT; the wild-type, thr4/THR4mc; the *cothr4* mutant expressing *CoTHR4*:*mCherry*.

**S5 Fig. Subcellular localization of CoThr4 during appressorium formation.**

Conidial suspensions of the *CoTHR4*:*mCherry*-complemented transformant were incubated on the coverslips at 24 °C for 15min, 2, 4, 6, and 24 h. Scale bar; 10 μm.

**S6 Fig. Conidia produced from threonine-treated *cothr4* mutant formed penetration hyphae on cellulose membranes in the presence of threonine.**

**(A)** Penetration hyphae of the *cothr4* mutant on the cellulose membranes. Scale bar; 10 μm. Conidia of each strain were suspended by 1 mM threonine and incubated on cellulose membranes at 24 °C for 48 h.

**(B)** The frequency of penetration hyphae of the *cothr4* mutant. Approximately 100 appressoria of each strain were observed, and two replicates were examined. Three independent experiments were conducted, and standard errors were indicated.

**S1 Table. PCR primers used in this study.**

